# DNA methylation reveals evolved buffering responses to climate-driven sex ratio skew in sea turtles

**DOI:** 10.64898/2026.01.06.697884

**Authors:** Eugenie C. Yen, Gabriella A. Carvajal, James O. Bazely, James D. Gilbert, Albert Taxonera, Kirsten Fairweather, Artur Lopes, Inês O. Afonso, Débora Newlands, Blair P. Bentley, Sandra M. Correia, Camryn D. Allen, Marylou K. Staman, Itzel Sifuentes-Romero, José M. Martín-Durán, Lisa M. Komoroske, Jeanette Wyneken, Christophe Eizaguirre

## Abstract

Species with temperature-dependent sex determination (TSD), including all sea turtles which produce females at warmer temperatures, face projections of demographic collapse under climate-driven sex ratio skews. However, the accuracy of such predictions remains uncertain, as current models rely heavily on indirect sex ratio proxies due to the lack of a scalable, non-invasive method for sexing hatchlings. Through whole methylome sequencing, we identified 777 sex-associated DNA methylation markers from blood samples of sex-verified loggerhead turtle (*Caretta caretta*) hatchlings incubated at three controlled temperatures. Applying these markers to a large-scale field experiment showed that classic nest temperature-based models overestimated female production by an average of up to 60.2%, suggesting the presence of evolved buffering mechanisms against thermal effects on sex determination. Our findings highlight the need to revise climate-driven sex ratio forecasts with empirical field data, such as methylation-based assessments, to better understand and safeguard the hidden resilience of vulnerable TSD species.

## Introduction

Climate change is rapidly reshaping ecosystems worldwide, with profound demographic consequences for broad taxa^1^. Over 400 species with temperature-dependent sex determination (TSD) are especially vulnerable, as embryonic sex is determined by temperature during a critical window of development, making them susceptible to extreme sex ratio skews as climate change intensifies^2^. These include all protected sea turtle species^3^, which develop as females at warmer incubation temperatures^4^. For many sea turtle populations, projections of near-complete female bias of hatchling sex ratios by the century end^5–9^ have fuelled concerns over climate-driven demographic collapse and extinction, potentially undermining decades of conservation success^10^.

However, the accuracy of these alarming forecasts remains uncertain. Current projections rely heavily on models that use nest temperature as a proxy for sex ratio, derived from limited constant-temperature laboratory experiments^11–13^. This method overlooks the complexity of natural systems, where temperatures fluctuate and microclimatic variation is high^14,15^. Moreover, the theoretical pivotal temperature (TPiv), which produces an equal sex ratio under a constant-temperature regime^16^, varies among populations, suggesting local adaptation of the TSD pattern to diverse thermal environments across the globe^17–19^. If heritable or plastic responses can buffer the thermal sensitivity of gonadal sex differentiation^20,21^, then current demographic forecasts may be substantially overestimating the magnitude of female bias. Large-scale empirical data on hatchling sex ratios are thus essential to evaluate whether adaptive mechanisms influence projections of demographic resilience, crucial for informing appropriate management decisions under climate change^22^.

Such progress towards empirical studies has been impeded by a fundamental methodological barrier, whereby hatchling sex cannot be identified easily due to the lack of heteromorphic sex chromosomes or sex-specific external morphology until maturity^23^. Existing methods for sexing hatchlings involve lethal sampling for gonadal histology, or several months of captive rearing before invasive gonadal laparoscopy, neither feasible at the scale necessary for comprehensive assessments^23^. Anti-Müllerian Hormone assays were initially promising, but have since proved irreproducible^24^. DNA methylation, the addition of a methyl group to cytosine bases, offers an alternative molecular solution for sexing hatchlings^25^. This epigenetic modification regulates gonadal TSD expression pathways^26,27^ and exhibits widespread sexual dimorphism across tissues and developmental stages^28^. If sex-diagnostic methylation markers from a peripheral tissue such as blood^29,30^ can be identified and even transferred between sea turtle populations, they would revolutionise our ability to directly assess hatchling sex ratios at scale in the field.

To address these longstanding fundamental and technical research gaps, we combined laboratory and field experiments at two of the world’s largest loggerhead turtle (*Caretta caretta,* IUCN Status: Vulnerable)^3^ nesting populations, together capturing most of the species’ global nesting distribution. We first identified methylation-based sex markers from minimally invasive blood samples of sex-verified Northwest Atlantic hatchlings, which were artificially incubated at three controlled temperatures. We next applied these novel markers to sex Northeast Atlantic hatchlings from a large-scale field experiment, where nests were exposed to natural thermal regimes. Finally, we compared methylation-based sex ratio predictions with classic nest temperature-derived estimates to evaluate how they impact assessments of demographic resilience under climate change.

## Results

### Identification of methylation-based sex markers

We used whole-genome bisulfite sequencing (WGBS) to profile nucleated red blood cell methylation of 29 sex-verified loggerhead turtle hatchlings from the Northwest Atlantic nesting population in Florida, Southeast USA (‘US-Inc’; **Fig. 1a**). To disentangle the effects of sex and temperature on methylation, these hatchlings were artificially incubated at male- (27°C), mixed- (30°C), and female-promoting (32°C) temperatures, with sex confirmed independently via laparoscopic examination of the gonads and their associated ducts^23^. Overall, 18 males (n=9 at 27°C, n=9 at 30°C) and 11 females (n=8 at 32°C, n=3 at 30°C) were sequenced across the three temperature treatments. From 4,816,224 cytosine-guanine dinucleotides (CpGs) covered in all hatchlings, we identified 777 differentially methylated sites (DMS: methylation difference >10%, q<0.05; **Extended Data Fig. 1**) between the sexes (n=383 female-hypermethylated, n=394 male-hypermethylated), with an absolute mean methylation difference of 21.3% across all DMS (range: 37.4% female-hypermethylated to 36.3% male-hypermethylated).

**Fig. 1.**
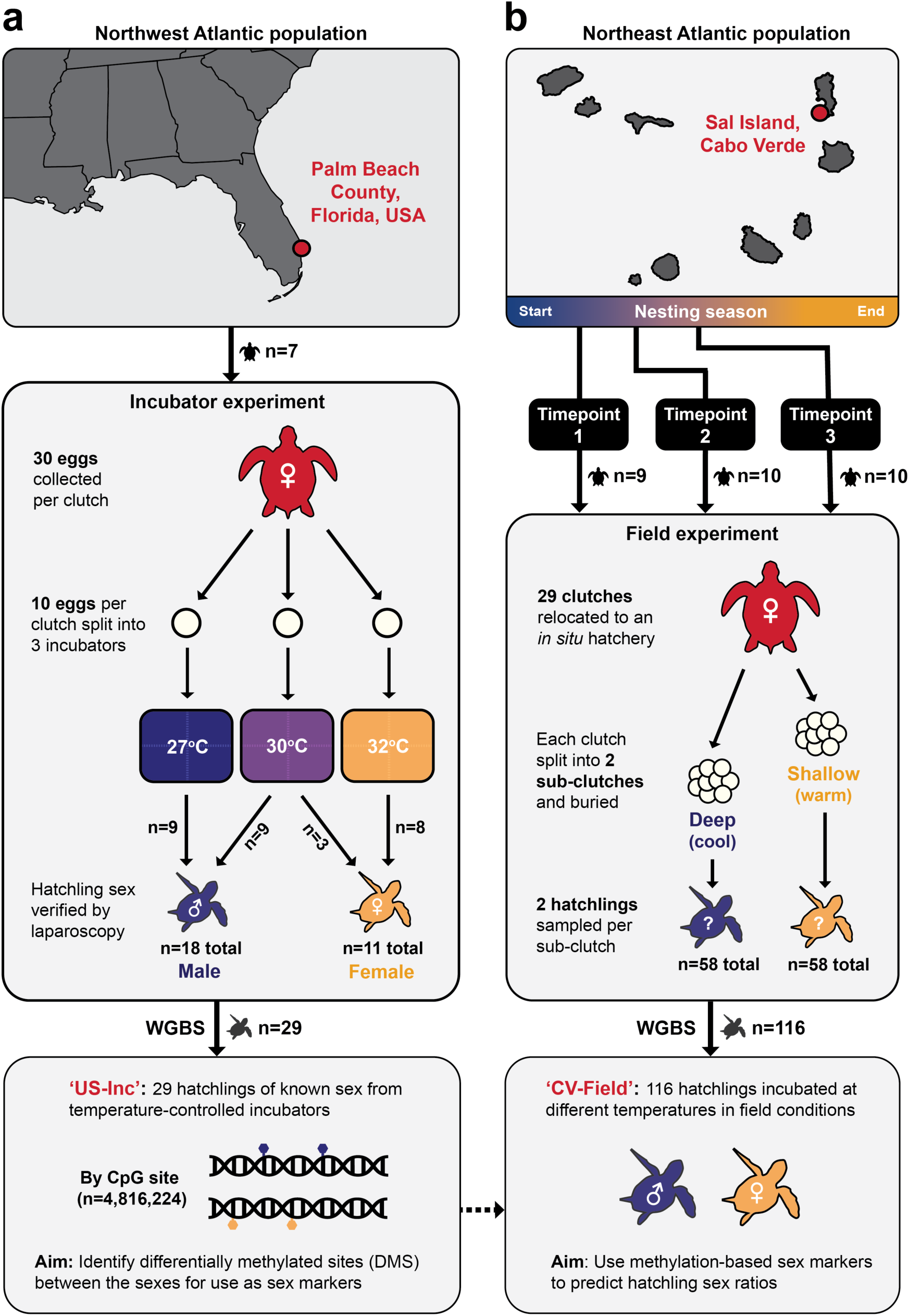
Experimental design. **(a)** Laboratory incubation experiment with the Northwest Atlantic loggerhead turtle nesting population. Eggs were collected from wild nesting females (n=7) then incubated under three controlled temperatures. Blood samples were collected from sex-verified hatchlings (‘US-Inc’, n=29) for whole genome bisulfite sequencing (WGBS). **(b)** Split-clutch field experiment with the Northeast Atlantic loggerhead turtle nesting population. Entire clutches of wild nesting females (n=29) were relocated to an *in situ* hatchery at three timepoints as temperatures rose over the first half of the nesting season (Timepoint 1: 9^th^ of July, Timepoint 2: 29^th^ of July, Timepoint 3: 23^rd^ of August). Each clutch was split and re-buried at cooler ‘deep’ and warmer ‘shallow’ treatments (n=58 sub-clutches). Blood samples were collected from hatchlings (‘CV-Field’, n=116) for WGBS.

Methylation variance at the 777 DMS clustered the US-Inc hatchlings strongly by sex via principal component analysis (PCA) and hierarchical clustering (**Figs. 2a, 2b**). Sex was confirmed as the sole factor associated with clustering (PERMANOVA: R²=0.46, F1,25=23.3, p=0.001), with no significant effect of temperature treatment or maternal ID (**Extended Data Table 1**). In contrast, global methylation variation at all 4,816,224 CpGs was associated with maternal ID (PERMANOVA: R^2^=0.04, F1,27=1.15, p=0.004; **Figs. 2c, 2d**), likely representing known genetic and maternal effects^31,32^. A weak association with temperature then emerged after accounting for intra-clutch relatedness (PERMANOVA: R^2^=0.04, F1,25=1.05, p=0.04). Altogether, these results confirm the successful isolation of sex-specific signals at the set of 777 DMS, independent of temperature or maternal effects observed at the global methylome level, and are thus primed for use as methylation-based sex markers.

**Fig. 2.**
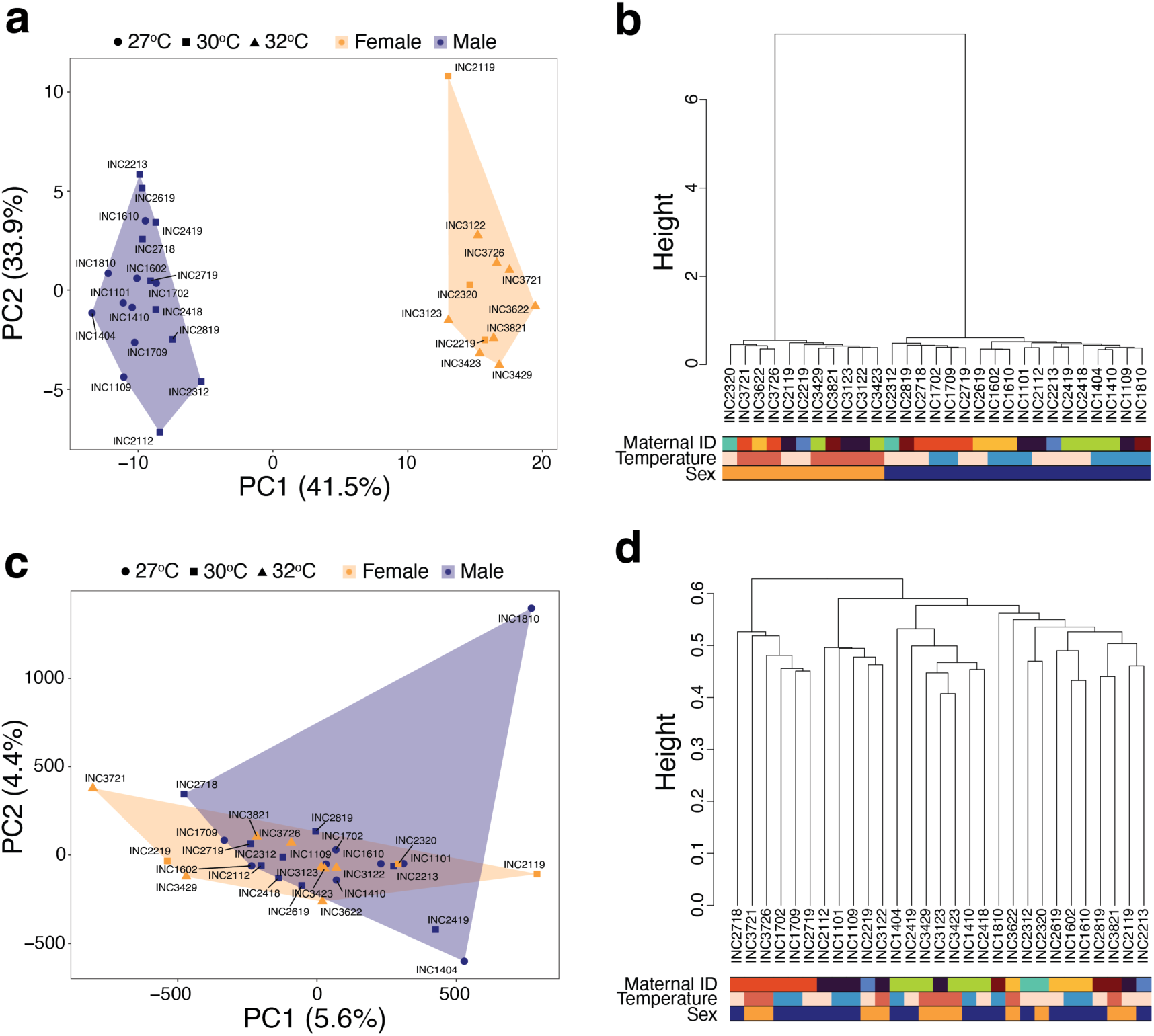
Methylation clustering of US-Inc hatchlings. **(a-b)** Methylation clustering of US-Inc hatchlings (n=29) at 777 differentially methylated sites (DMS) between the sexes. **(a)** Principal component analysis (PCA), with the proportion of methylation variance explained by the top two principal components (PCs) in brackets on each axis. Each point is a hatchling ID, where colour represents sex (females: yellow, males: blue) and shape represents temperature treatment (27°C: circle, 30°C: square, 32°C: triangle). **(b)** Hierarchical clustering dendrogram, where each branch is a hatchling ID. For each hatchling, the coloured top bar represents maternal ID, the middle bar represents temperature treatment (27°C: blue, 30°C: peach, 32°C: red), and the bottom bar represents sex (females: yellow, males: blue). **(c-d)** Global methylation clustering of US-Inc hatchlings (n=29) at all 4,816,224 genome-wide CpGs, where **(c)** is a PCA and **(d)** is a hierarchical clustering dendrogram, following the same specifications as described for **(a)** and **(b)**, respectively.

### Functional context of methylation-based sex markers

A total of 436 DMS were associated with 392 annotated genes by overlap with genic feature types (promoter: n=21, exon: n=12, intron: n=331), or proximity to a transcriptional start site (intergenic: n=72; **Supplementary Table 1**). Gene Ontology analysis revealed diverse functional enrichments in DMS-associated genes, including metabolic regulation, cell signalling, and cellular transport **(Extended Data Fig. 2, Supplementary Table 2).** Notably, from peripheral blood samples, we still captured persistent differential methylation at 40 genes with documented roles in sex development or gametogenesis (**Fig. 3a, Supplementary Table 3**), despite the general tissue- and timing-specific nature of methylation patterning^33^. For instance, the spermatogenesis-related CAMK4 gene^34,35^ had six male-hypermethylated DMS (mean methylation difference: 20.7%, intronic; **Fig. 3b**). The most female-hypermethylated DMS was on the AMDHD2 gene promoter (difference: 34.5%; **Fig. 3b**) and is implicated in metabolism during spermatogenesis^36^, whereas the most male-hypermethylated DMS was associated with HSP90AB1 (difference: 27.8%, intergenic; **Fig. 3b**) and has been linked to the TSD gonadal cascade^37^. Beyond sex development genes, we identified the top 5% methylation difference outliers overall (n=29 DMS, n=14 DMS-associated genes; **Fig. 3a, Extended Data Table 2**). The TRIT1 gene, with no known sex-related functions, had four outlier DMS (mean difference: 32.2%, intronic; **Fig. 3b**), including the most female-hypermethylated DMS (difference: 37.4%). Meanwhile, the most male-hypermethylated DMS was associated with PCDH10 (difference: 32.9%, intergenic; **Fig. 3b**), a neurodevelopmental gene involved in sex-dependent fear responses^38^. These results demonstrate that methylation sex differences detectable from hatchling blood also reflect systemic sexual dimorphism of broad tissues extending beyond the sex organs.

**Fig. 3.**
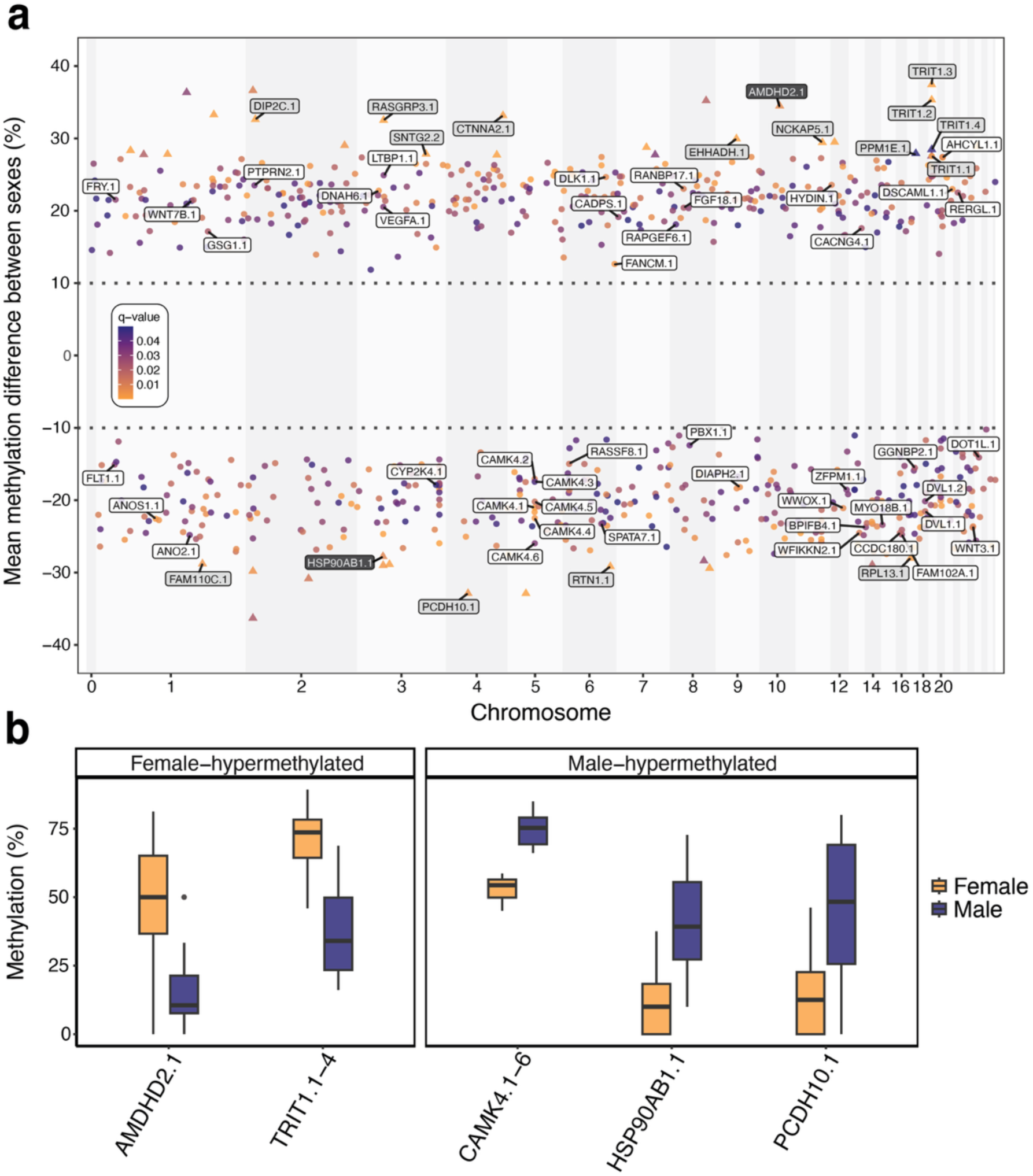
Genes underlying sex-specific methylation in US-Inc hatchlings. (**a**) Manhattan plot of 777 differentially methylated sites (DMS) between the sexes of US-Inc hatchlings. DMS are plotted along the 28 chromosomes of the loggerhead turtle genome (chromosome 0: unplaced contigs), against the mean methylation difference (%) between sexes (positive: hypermethylated in females, negative: hypermethylated in males). Points are coloured by q-value and triangles represent DMS that fell within the top 5% outlier subset (n=29) in relation to absolute mean methylation difference between the sexes. Labelled gene IDs indicate genes with documented roles in sex development or gametogenesis (white), genes with DMS within the top 5% outlier subset (grey), and genes that fulfilled both categories (black). (**b**) Mean methylation (%) per hatchling (n=29) for select DMS, coloured by sex (females: yellow, males: blue). For all boxplots, the central line represents the median, the box depicts the interquartile range (IQR), and the whiskers represent 1.5 times the IQR. From the left, AMDHD2.1 is the most female-hypermethylated DMS on a sex development-related gene; TRIT1.1-4 shows the mean across 4 DMS on the TRIT1 gene, which hosted the most female-hypermethylated DMS overall and also the largest number of DMS within the top 5% outlier subset; CAMK4.1-6 represents the mean across 6 DMS on the CAMK4 gene, which had the largest number of DMS overall for a sex development-related gene; HSP90AB1.1 is the most male-hypermethylated DMS on a sex development-related gene; PCDH10.1 is the most male-hypermethylated DMS overall.

Gene expression regulation was not correlated with methylation levels for 178 DMS-associated genes (45.4%) captured in paired RNA-Seq data from the same blood samples (Wald χ^2^=0.15, p=0.70; methylation x feature type: Wald χ^2^=0.028, p=0.99, n=18 hatchlings) (Carvajal *et al., in prep*). This supports that sex-associated methylation signatures at the DMS represent stable epigenetic marks, rather than solely reflecting current transcriptional activity of blood cells.

### Application of methylation-based sex markers in the field

To test our methylation-based sex markers on hatchlings incubated under natural thermal regimes, we conducted a split-clutch field experiment at the Northeast Atlantic nesting population in Cabo Verde (CV-Field; **Fig. 1b**). We relocated 29 wild clutches (81.3 ± 12.3 eggs) to an *in situ* hatchery at three timepoints over the warming nesting season. Each clutch was split in half (n=58 sub-clutches) and re-buried at a cooler ‘deep’ (55 cm) treatment and warmer ‘shallow’ (35 cm) treatment, thus controlling for maternal effects, while inducing thermal variation^32^. From temperature logger data for 34 sub-clutches (**Fig. 4a**), we verified that the experimental design had induced different incubation temperatures. Mean temperature per timepoint-treatment (n=6 combinations) spanned 29.8°C to 31.7°C (**Table 1**, **Fig. 4b**), while the overall temperature range was 27.2–34.2°C (**Supplementary Table 4**). Shallow sub-clutches were consistently 0.7°C warmer than deep sub-clutches (ANOVA: F1,30=109.7 p<0.0001), with mean temperature also rising across successive timepoints (Timepoint 1 to 2: +0.9°C, Timepoint 2 to 3: +0.4°C; ANOVA: F1,30=76.3, p<0.0001).

**Fig. 4.**
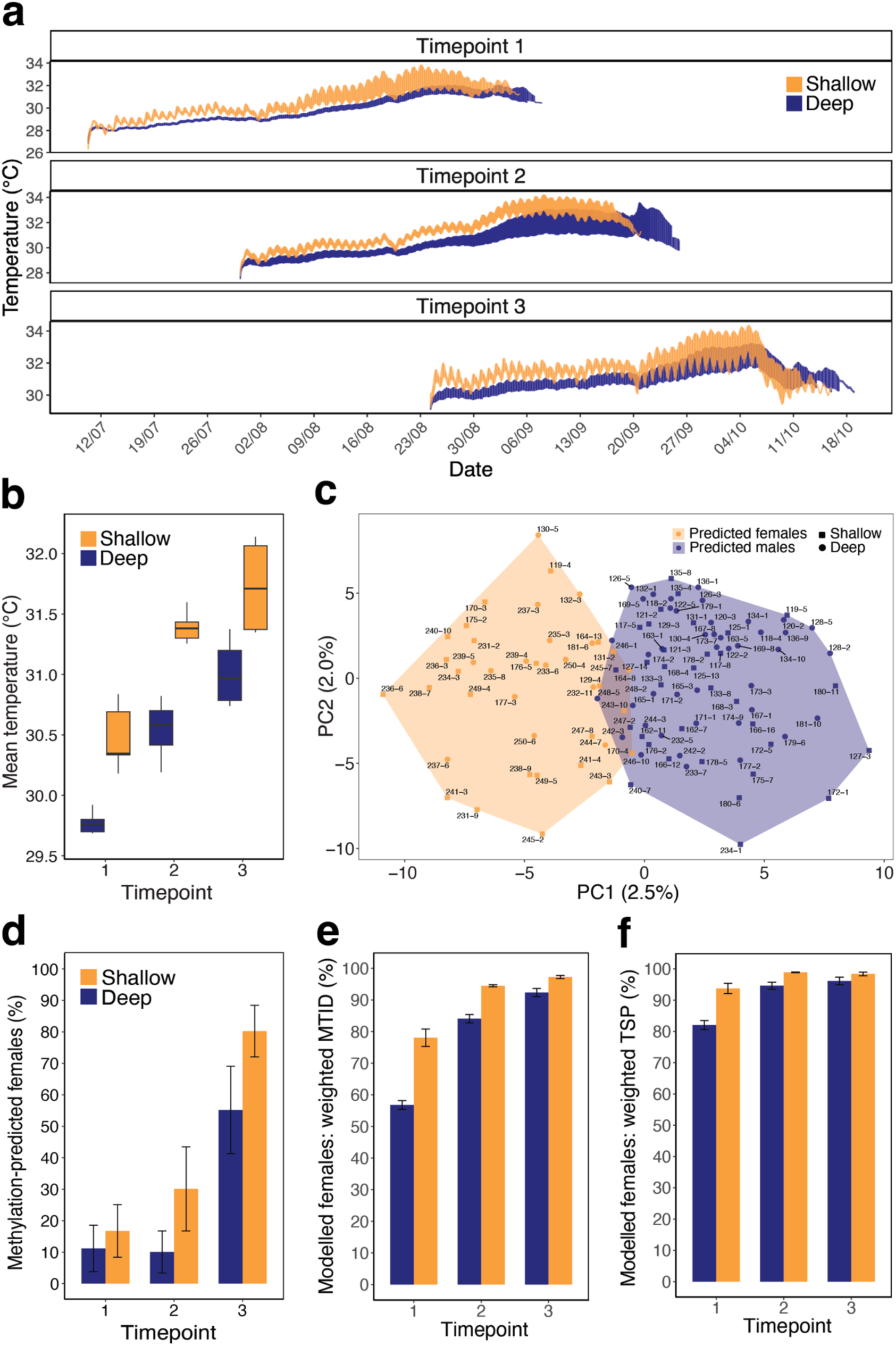
CV-Field hatchling sex ratios under natural thermal regimes. **(a)** Temperature series (°C) over the full incubation period for 34 sub-clutches (Timepoint 1: n=5 per treatment, Timepoint 2: n=8 per treatment, Timepoint 3: n=4 per treatment) of the field experiment, coloured by depth treatment (shallow: yellow, deep: blue). **(b)** Mean temperature (°C) during the full incubation period across sub-clutches with temperature data (n=34 total) per timepoint and depth treatment (shallow: yellow, deep: blue). The central line represents the median, the box depicts the interquartile range (IQR), and the whiskers represent 1.5 times the IQR. **(c)** Principal component analysis of CV-Field hatchlings (n=116) at 743 CpGs that overlapped with DMS identified from US-Inc hatchlings. The proportion of methylation variance explained by the top two principal components (PCs) is shown in brackets on each axis. Each point is a hatchling ID, where colour represents methylation-predicted sex clusters (predicted females: yellow, predicted males: blue), and shapes represent depth treatments (shallow: square, deep: circle). **(d-f)** Mean hatchling sex ratio estimates per timepoint and depth treatment. For all bar charts, colour represents depth treatment (shallow: yellow, deep: blue) and error bars are standard errors. **(d)** Mean methylation-predicted sex ratios, calculated across all CV-Field hatchlings (Timepoint 1: n=18 hatchlings per treatment; Timepoints 2 and 3; n=20 hatchlings per treatment; n=116 total). **(e-f)** Mean modelled sex ratios, calculated across all sub-clutches with temperature series data (n=34 total) during **(e)** the weighted middle third of incubation (MTID), and **(f)** the weighted thermosensitive period (TSP).

**Table 1.**
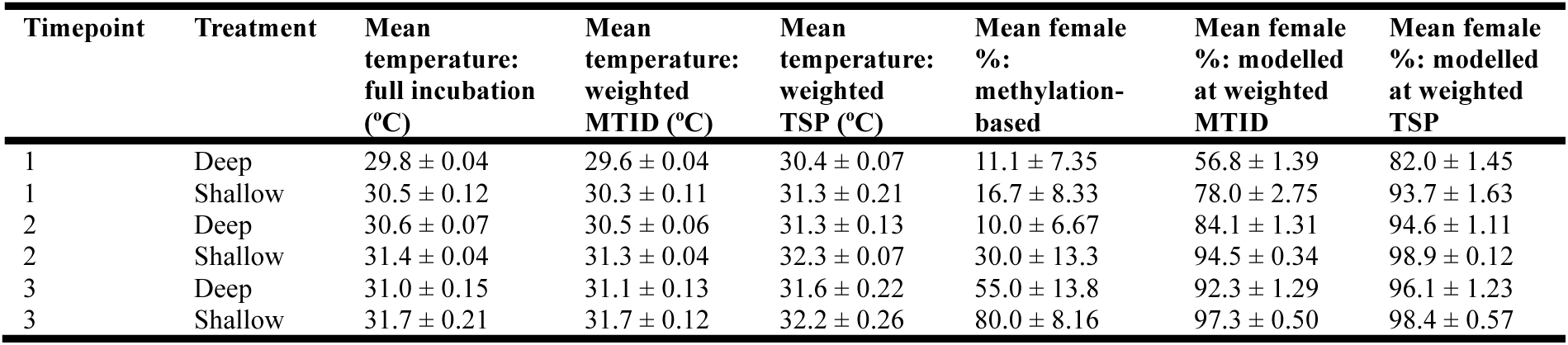
Incubation temperatures and sex ratios of CV-Field hatchlings. For each timepoint and depth treatment in the field experiment, the mean temperature (°C ± standard error, SE) is shown across all sub-clutches with available temperature data (Timepoint 1: n=5 per treatment, Timepoint 2: n=8 per treatment, Timepoint 3: n=4 per treatment; n=34 total). These were calculated for the full incubation period, as well as constant temperature equivalents derived from a thermal embryonic growth model during the weighted middle third of incubation (MTID) and weighted thermosensitive period (TSP). Mean methylation-predicted female proportions (% ± SE) were calculated across all CV-Field hatchlings per timepoint-treatment (Timepoint 1: n=18 hatchlings per treatment, Timepoints 2 and 3: n=20 hatchlings per treatment; n=116 total). Mean modelled female proportions (% ± SE) were calculated across all sub-clutches with temperature series data (n=34 total) during the weighted MTID and weighted TSP.

| Timepoint | Treatment | Mean temperature: full incubation (°C) | Mean temperature: weighted MTID (°C) | Mean temperature: weighted TSP (°C) | Mean female %: methylation-based | Mean female %: modelled at weighted MTID | Mean female %: modelled at weighted TSP |
| --- | --- | --- | --- | --- | --- | --- | --- |
| 1 | Deep | 29.8 ± 0.04 | 29.6 ± 0.04 | 30.4 ± 0.07 | 11.1 ± 7.35 | 56.8 ± 1.39 | 82.0 ± 1.45 |
| 1 | Shallow | 30.5 ± 0.12 | 30.3 ± 0.11 | 31.3 ± 0.21 | 16.7 ± 8.33 | 78.0 ± 2.75 | 93.7 ± 1.63 |
| 2 | Deep | 30.6 ± 0.07 | 30.5 ± 0.06 | 31.3 ± 0.13 | 10.0 ± 6.67 | 84.1 ± 1.31 | 94.6 ± 1.11 |
| 2 | Shallow | 31.4 ± 0.04 | 31.3 ± 0.04 | 32.3 ± 0.07 | 30.0 ± 13.3 | 94.5 ± 0.34 | 98.9 ± 0.12 |
| 3 | Deep | 31.0 ± 0.15 | 31.1 ± 0.13 | 31.6 ± 0.22 | 55.0 ± 13.8 | 92.3 ± 1.29 | 96.1 ± 1.23 |
| 3 | Shallow | 31.7 ± 0.21 | 31.7 ± 0.12 | 32.2 ± 0.26 | 80.0 ± 8.16 | 97.3 ± 0.50 | 98.4 ± 0.57 |

We next performed WGBS on blood samples from two hatchlings per sub-clutch (n=116 total, Timepoint 1: n=18 per treatment, Timepoints 2 and 3: n=20 per treatment). Of 777 DMS identified between the sexes of US-Inc hatchlings, 743 (95.6%) were covered in the CV-Field hatchling dataset and hence applied as methylation-based sex markers. K-means clustering at these 743 sites optimally revealed two discrete natural clusters (**Extended Data Fig. 3a**), independent of timepoints or depth treatments (**Fig. 4c**). We assigned cluster identity to predicted sex based on two expectations under the sea turtle TSD system: the proportion of females should (1) increase as temperatures rose across timepoints, and (2) be consistently higher in warmer shallow treatments than cooler deep treatments within each timepoint. The proportion of Cluster 1 hatchlings followed both expectations precisely, increasing over timepoints (Timepoint 1: 13.9%, Timepoint 2: 20%, Timepoint 3: 67.5%) with more shallow-incubated hatchlings per timepoint (Timepoint 1shallow-deep: 5.6%, Timepoint 2shallow-deep: 20%, Timepoint 3shallow-deep: 25%; **Extended Data Fig. 3b**). Hence, natural clustering patterns supported the assignment of Cluster 1 as predicted females and Cluster 2 as predicted males.

Methylation clustering of the CV-Field hatchlings was associated with predicted sex cluster identity as expected (PERMANOVA: R^2^=0.020, F1,108=2.33, p=0.001), alongside effects of maternal ID (PERMANOVA: R^2^=0.26, F1,108=1.12, p=0.001) and depth treatment (PERMANOVA, R^2^=0.010, F1,108=1.23, p=0.001; **Extended Data Table 3**). A combined PCA of US-Inc and CV-Field hatchlings also revealed that the sexes separated along different principal components between populations (**Extended Data Fig. 3c**). Together, these results indicate that methylation-based sex markers are generally transferable across Atlantic populations, while simultaneously capturing population-specific molecular signatures.

### Comparison of methylation-based and temperature-modelled sex ratios

Methylation-based predictions yielded 40 females out of 116 CV-Field hatchlings (34.5%) in the field experiment. The mean female proportion per timepoint-treatment ranged from 11.1% to 80.0%, with early male bias at Timepoints 1 and 2 shifting to female bias by Timepoint 3 (**Table 1, Fig. 4d**). For comparison, we modelled sex ratios using a well-established nest temperature-based method^11,13^. For 34 sub-clutches with temperature data, we derived the constant temperature equivalent during two standard proxies for the sex determining window: (1) the middle third of incubation duration weighted by embryo growth rate (weighted MTID), and (2) the thermosensitive period (middle third of total development) weighted by embryo growth rate and the thermal reaction norm of sexualization (weighted TSP)^13^. Sex ratios were then estimated using the constant-temperature TSD pattern for a Northwest Atlantic loggerhead population^39^, with a TPiv of 29.44°C, as this parameter remains unvalidated for Cabo Verde. Both models consistently predicted high female bias throughout the experiment, with the proportion of females per timepoint-treatment ranging between 56.8–80.0% at the weighted MTID (mean: 83.8%; **Fig. 4e**) and 82.0–98.4% at the weighted TSP (mean: 94.0%; **Fig. 4f**, **Table 1, Extended Data Fig. 4**). Such variation in model estimates derived from the same temperature series data emphasises the need for empirical validation.

When compared to methylation-based predictions, we showed that temperature-based modelling overestimated female hatchling production by an average of 50.0% at the weighted MTID and 60.2% at the weighted TSP, with maximum overestimations of 74.1% and 84.6%, respectively. Of note, Timepoint 3 preceded peak nesting activity in the experimental year, with 36,952 out of 53,498 (69.2%) clutches laid after this point (**Extended Data Fig. 5**). Therefore, though less extreme than temperature-modelled estimates, methylation-based predictions still indicate overall female hatchling bias at the population level, as most nests will experience warmer temperatures over the remainder of the nesting season. Sex ratios were further estimated from half-sized clutches (40.2 ± 6.2 eggs) in this experiment, so natural clutches are also expected to produce more females under stronger metabolic heating effects^40^.

## Discussion

Species with temperature-dependent sex determination (TSD) may face extremely skewed sex ratios as climate change progresses, threatening their long-term persistence. Yet, evaluating this risk in sea turtles has been constrained for decades by the lack of a scalable, non-invasive method for sexing hatchlings. Here, we experimentally identified DNA methylation markers from blood samples that distinguish hatchling sex in loggerhead turtles, then applied them to measure hatchling sex ratios at a globally important Northeast Atlantic nesting population. The stark mismatch between our empirical sex ratio predictions and nest temperature-derived models exposes fundamental uncertainties in current climate vulnerability assessments. This finding has immediate implications for forecasting and managing climate impacts on TSD species.

From a controlled experiment with Northwest Atlantic hatchlings, we identified 777 methylation-based markers that successfully isolated sex-specific signals from confounding thermal and genetic variation observed at the global methylome level. This underscores the strength of a methylome-wide discovery approach for identifying reliable correlates of sex in wild species, building upon earlier efforts in alligators^29^. The functional context of these markers strengthens confidence in their biological relevance. Despite profiling peripheral blood rather than gonadal tissue, we captured persistent differential methylation at 40 genes with documented roles in sex development or gametogenesis, including AMDHD2, which independently emerged as a sex-specific marker in humans and thus has promising cross-species applicability^41^. Approximately 45% of marker-associated genes exhibited transcriptional activity, indicating that methylation signatures can reflect stable epigenetic marks maintained across tissues and developmental stages, beyond transient transcriptional states^28,42^. This systemic stability supports reliable marker identification and confers practical advantages to DNA methylation over expression-based approaches, where distinguishing true biological absence from detection limits remains challenging. DNA methylation also remains stable across diverse storage conditions^43^, enhancing its logistical feasibility for application in remote field settings.

When applied to Northeast Atlantic hatchlings incubated under natural thermal regimes, traditional temperature-based models overestimated female hatchling production by an average of 50.0% or 60.2% compared to empirical methylation-based predictions. This systematic overestimation likely reflects two compounding limitations on model performance: the inherent inadequacy of laboratory-derived, constant-temperature parameters when applied to fluctuating field conditions^14,15,43^, and the reliance on a non-local pivotal temperature (TPiv: 29.44°C)^38^. Directly transposing TPiv in this manner is standard practice when local validation is unavailable, which remains the case for most populations, yet is problematic given evidence for inter-population variation in TPiv17,18. Previous sex ratio projections for the Cabo Verde population used an even lower TPiv of 29°C^6,8^, suggesting published estimates of climate-driven sex ratio bias may be substantially inflated. Meanwhile, our empirical approach was independently validated by the close alignment of methylation-based sex ratio predictions with expected seasonal and thermal trends in our field experiment. Unlike the models, these trends emerged naturally without being guided by environmental parameters. Methylation-based sexing further yields individual-level sex phenotypes, not just average hatchling sex ratios.

All else being equal, our methylation-based results suggest a field-equivalent TPiv above 31°C, since we observed both 30% and 55% female hatchling proportions over this temperature during the incubation period. While a TPiv of this magnitude has been documented in other sea turtle species^18^, it exceeds previous estimates for loggerhead turtles^4,39^. This higher TPiv has striking implications for the demographic resilience of sea turtles under climate change, as it suggests that some populations have a greater capacity to buffer the developmental outcomes of sex determination against thermal variation than assumed by current models^6,8^. As such, our study exposes a form of hidden resilience, defined here as molecular or physiological flexibility that alters the thermal sensitivity of gonadal sex differentiation beyond what laboratory-derived parameters predict. This interpretation is consistent with adaptation of populations to local environmental regimes, where selection favours the evolution of buffering mechanisms against extreme sex ratio skews, due to the reproductive fitness advantage experienced by the rarer sex^45,46^. The difference in methylation profiles we observed between the Northwest and Northeast Atlantic populations, which separated along different principal components despite sharing core sex-associated patterns, provides early molecular support for such divergence.

Importantly, we do not suggest that climate change poses no demographic risk to sea turtle persistence. While less extreme than previous projections, our methylation-based estimates still indicate female-biased hatchling sex ratios for the Northeast Atlantic population. Climate warming therefore remains a pressing concern requiring active monitoring and management, although the disparity in magnitude of resilience fundamentally alters the urgency of climate vulnerability assessments. Deciphering whether the molecular basis of this hidden resilience is heritable and/or plastic will be essential for predicting whether the adaptive potential of sea turtles can match the contemporary pace and intensity of climate change^47^. Addressing this gap will require large-scale empirical studies that quantify how population-specific adaptive responses impact true hatchling sex ratio outcomes. Methylation-based approaches can play a central role in achieving this goal, as the top sex-diagnostic markers can be upscaled via targeted methylation assays for widespread application across wild species and populations^25^.

By facilitating scalable, non-invasive assessments of hatchling sex ratios in the field, DNA methylation offers a promising empirical foundation to improve the realism of climate risk projections for long-term conservation planning. It further paves the way for precision conservation approaches, where molecular markers are deployed in real-time to monitor at-risk populations and evaluate the success of interventions such as nest shading, relocation, and assisted adaptation. More broadly, our findings illustrate that molecular approaches can uncover unexpected forms of resilience, even in the most climate-vulnerable species. Understanding and harnessing this hidden resilience will be essential for safeguarding TSD species effectively in the Anthropocene.

## ONLINE METHODS

### Incubator experimental design (US-Inc)

We set up a laboratory-based, temperature-controlled incubator experiment using eggs sampled from the main Northwest Atlantic loggerhead nesting population (Florida, USA; **Fig. 1a**). During the July 2022 nesting season, 30 eggs each from seven wild clutches (240 eggs total) were collected the morning after oviposition at Boca Raton (26.60°N, -80.10°E) and Juno (26.70°N, -80.05°E) Beaches in Palm Beach County, Florida, USA. Eggs were transported for 1-2 hours in vacuum-sealed bags to a laboratory at Florida Atlantic University (Florida, USA). Here, artificial incubators were maintained at three controlled temperatures to disentangle temperature from sex effects on observed methylation patterns: 27°C (male-promoting temperature), 30°C (mixed sex-promoting temperature), and 32°C (female-promoting temperature). Each incubator housed four Styrofoam^®^ (Dow Chemical Company, USA) nest boxes filled with sterilized beach sand, with eggs from each clutch distributed evenly and quasi-randomly among nest boxes to control for maternal effects^24^. Temperature was continuously monitored at 15 min intervals using HOBO U22 (Onset Computer Corporation, USA) dataloggers (± 0.2°C accuracy)^15^. To control for sand moisture effects, the volumetric moisture content was measured daily by oven-drying and weighing 40–60 g of sand per nest box, targeting an average moisture of 6% (0.06 m³/m³), consistent with natural nests^48^.

Blood samples were collected from 1-3-day old hatchlings, once the plastron was flat and the egg yolk sac was fully internalised. A maximum of 7% of the total body mass in blood volume was collected per hatchling^49^ from the external jugular vein with a 26-gauge half-inch needle and 1 mL heparinised syringe^50^. Samples were centrifuged for 1-2 min at 6000 rpm to separate plasma and nucleated erythrocytes, then stored at -80°C. Hatchlings were transported to Florida Atlantic University Marine Science Laboratory (Florida, USA) and captive reared for 2.5-6 months until they were large enough for sex verification via laparoscopic examination of the gonads and their associated ducts^23^. A total of 29 hatchlings were selected for WGBS (**Fig. 1a**; ‘US-Inc’ hatchlings), spanning the three temperature treatments and both sexes: 18 males (n=9 at 27°C, n=9 at 30°C) and 11 females (n=8 at 32°C, n=3 at 30°C), across the seven clutches (**Supplementary Table 5**). Genomic DNA was extracted from the nucleated red blood cell samples with a Monarch Spin gDNA Extraction Kit (New England Biolabs, USA) following the manufacturer’s protocol. WGBS libraries (Novogene, USA) were constructed and sequenced with 150 bp paired-end reads on an Illumina NovaSeq 6000 platform (Illumina, USA). To supplement low initial depth of coverage, a second round of DNBseq WGBS libraries were constructed from the same DNA extraction batch and sequenced with 100 bp paired-end reads on an MGI DNBSEQ platform (MGI Tech, China).

### Field experimental design (CV-Field)

We conducted a field experiment at the main Northeast Atlantic loggerhead turtle nesting population (Cabo Verde; **Fig. 1b**) during the 2021 nesting season. Immediately after oviposition, we collected 29 entire clutches (81.3 ± 12.3 (standard deviation, SD) eggs) from wild turtles that nested on Algodoeiro Beach (16.62123°N, -22.92972°E), Sal Island, Cabo Verde over three nights: Timepoint 1 on the 9^th^ of July (n=9 clutches), Timepoint 2 on the 29^th^ of July (n=10 clutches), and Timepoint 3 on the 23^rd^ of August (n=10 clutches). This design generated a range of natural nest incubation temperatures as ambient temperatures rose over the experimental nesting season period, while controlling for inter-clutch variation per relocation date.

Within six hours of collection, clutches were relocated to a protected hatchery enclosure on a nesting beach (Project Biodiversity, Cabo Verde). There, we set up a split-clutch experiment to induce further variation in incubation temperature, while controlling for genetic and maternal effects^32^. Each clutch was randomly split into two sub-clutches of equal size to reduce metabolic heating effects (n=58 sub-clutches total; 40.2 ± 6.2 eggs). One sub-clutch was re-buried with a bottom egg chamber depth of 55cm as the cooler, ‘deep’ treatment, and the other at 35cm as the warmer, ‘shallow’ treatment. Both depths lie within the natural range observed for wild loggerhead turtle nests in Cabo Verde (38–67 cm)^51^. A HOBO Pendant MX Water Temp™ (Onset Computer Corporation, USA) logger was placed at the centre of 34 sub-clutches to record temperature at 15 min intervals (± 0.5°C accuracy), with paired loggers per depth treatment from the same maternal ID (Timepoint 1: n=5 loggers per treatment, Timepoint 2: n=8 loggers per treatment, Timepoint 3: n=4 loggers per treatment; **Supplementary Table 6**).

Upon natural emergence, we collected 50-100 μl of blood from the dorsal cervical sinus of each hatchling with a 26-gauge needle and 1 mL syringe. Samples were stored immediately in lithium heparin-coated tubes, then centrifuged for 1 min at 3000 rpm to separate plasma and red blood cells within 12 hours of collection. These were stored at -18°C until the field season ended, then at -80°C at Queen Mary University of London (UK). We sequenced red blood cell samples from two hatchlings per deep and shallow sub-clutch (**Fig. 1b**; ‘CV-Field’ hatchlings), selected following the logic outlined in Yen *et al.,* (2024). This resulted in 116 hatchlings sequenced over 58 sub-clutches (Timepoint 1: n=18 hatchlings per treatment; Timepoints 2 and 3: n=20 hatchlings per treatment). Genomic DNA was extracted with a DNeasy^®^ Blood and Tissue Kit (Qiagen, Germany) following the manufacturer’s protocol for nucleated red blood cells. DNBseq WGBS libraries were constructed and sequenced with 100 bp paired-end reads on an MGI DNBSEQ platform (MGI Tech, China).

### Methylation data processing

Summary statistics for each processing step are provided in **Supplementary Table 5** (US-Inc hatchlings) and **Supplementary Table 6** (CV-Field hatchlings). Raw WGBS reads were trimmed using cutadapt v.2.10^52^ to remove adapters and reads of Phred score <Q20. Trimmed reads were aligned against the chromosome-level reference genome assembly for a Northeast Atlantic loggerhead turtle^53^ using Bismark v.0.22.1 with default options^54^. Alignments were deduplicated in paired end mode with Bismark, then merged and sorted with SAMtools v.1.9^55^. For the US-Inc hatchlings, we performed quality control, alignment, deduplication, and sorting separately per batch to account for possible sequencing batch effects, before merging the datasets per CpG site with SAMtools^55^. This resulted in a total of 169,092,377 ± 29,516,897 reads per merged US-Inc sample and 134,716,463 ± 3,699,824 reads per CV-Field sample.

Methylation calling at CpGs was conducted via Bismark^54^. Bisulfite conversion efficiency estimated by totalling percentage methylation across CHG and CHH sites (US-Inc range: 96.5-97%, CV-Field range: 98.9-99.6%)^56^. As methylation occurs symmetrically across CpGs in vertebrates, CpGs were destranded to minimise pseudo-replication of cytosine methylation counts. This yielded 25,541,603 ± 139,468 CpGs with a destranded depth of coverage of 9.14 ± 1.37X per US-Inc hatchling, and 25,182,966 ± 644,316 CpGs with a destranded depth of coverage of 8.91 ± 0.84X per CV-Field hatchling. Methylation calls were processed in RStudio v.4.2.257 using methylKit v.1.32.1^58^. CpGs were filtered for a depth of coverage between 5X and the 99.9^th^ percentile to minimise PCR bias^59^, then normalised across samples within each dataset.

### Identification of methylation-based sex markers

To identify robust methylation-based sex markers, we applied a stringent filter that retained CpGs covered in all 29 US-Inc hatchlings, leaving 4,816,224 CpGs. We tested for differential methylation between the sexes using PQLseq v.1.2.1 with default parameters^60^, which fit a binomial mixed effects model per CpG, while accounting for genetic covariance by including a kinship matrix as a random effect. The kinship matrix set within-clutch relatedness to r=0.5 and between-clutch relatedness to r=0^32,61^. Non-converged sites were excluded and the sliding linear model method was applied to correct for multiple testing^62^. Mean methylation difference (%) between sexes per CpG was calculated with methylKit’s ‘calculateDiffMeth’ function as the difference between female minus male hatchlings, weighted by read coverage^58^. A CpG was classified as a differentially methylated site (DMS) if it had a q-value <0.05 and an absolute mean methylation difference >10% between the sexes^32^.

To verify that the DMS effectively separated hatchlings by sex without an influence of temperature or maternal effects, we performed a PCA using ‘prcomp’^57^ and hierarchical clustering in methylKit via Ward agglomeration with the correlation distance method. We tested for statistical associations with methylation clustering via PERMANOVA analyses with a Bray-Curtis distance matrix of methylation level (%) at the 777 identified DMS for all 29 US-Inc hatchlings. This was implemented via ‘adonis2’ in vegan v.2.6.4^63^ with 999 permutations. An initial PERMANOVA model tested maternal ID as the sole fixed effect on methylation clustering. If significant, maternal ID was incorporated as a stratification factor to account for intra-clutch non-independence in a subsequent PERMANOVA model examining the fixed effects of sex, temperature treatment, and their interaction on methylation clustering. To understand clustering at the global methylation level, we repeated the PCA, hierarchical clustering, and PERMANOVA analyses for all 4,816,224 CpGs covered in the genome.

### Characterisation of genes underlying methylation-based sex markers

Gene annotations^53^ were assigned using genomation v.1.30.0^64^ and GenomicRanges v.1.50.2^65^. Promoter regions were defined as 1500 bp upstream and 500 bp downstream of the transcriptional start site (TSS)^66^, then each CpG was classified into one of four feature types (promoter, exon, intron or intergenic). A CpG was considered to be gene-associated if it overlapped genic feature types or was located within 10 kbp of the nearest TSS when intergenic^66^. For genes lacking functional annotations in the reference genome^53^, BLASTp searches were conducted via NCBI’s web portal^67^ against the ClusteredNR database to identify potential homology-based functions (accessed 13^th^ June 2025). Hits were considered strong if they met the criteria: e-value <10^-30^, percentage identity >70%, percentage query cover >70%. This added functional gene information for 10 out of 57 previously unannotated genes (**Supplementary Table 1**).

A conditional hypergeometric Gene Ontology (GO) term enrichment analysis was performed with GOstats v.64.0^68^ and GSEABase v.1.60.0^69^. Gene sub-universes comprised of all DMS-associated genes with available GO annotations (n=206), split into genes hyper-methylated in females or males. These were compared against the gene universe of all genes associated with covered CpGs in the genome (n=13,377). Overrepresented GO terms were identified with the Benjamini-Hochberg method at a false discovery rate of 0.05^70^.

To identify genes with sex differentiation and development functions, we conducted a targeted literature search using Google Scholar (accessed 4^th^ September 2024). For each DMS-associated gene, Boolean search queries matched the gene ID with “sex” as keywords. Top results were manually screened for evidence of roles in sex organ development and/or gametogenesis. In addition, we cross-referenced DMS-associated genes against 223 genes with documented roles in TSD^37^. Finally, we characterised the top 5% methylation difference outliers across all DMS.

### Correlation between DNA methylation and gene expression

To explore the relationship between DNA methylation and expression levels at DMS-associated genes, we analysed RNA-Seq data available for 18 US-Inc hatchlings from Carvajal *et al.,* (*in prep*). Detailed RNA processing methods are reported in Carvajal *et al.,* (*in prep*). Briefly, RNA was extracted and sequenced from the same blood samples used for WGBS in this study. Reads were mapped to the ‘GSC_CCare_1.0’ loggerhead turtle reference genome^71^ using STAR v.2.7.11b^72^, then gene counts were produced using HTSeq v.2.0.4 with the ‘gene_id’ and ‘reverse’ stranded parameters^73^. Out of 392 DMS-associated genes, 307 shared functional gene names with the expression dataset. To ensure orthologous genes were identified between reference genomes, we retained genes on the same chromosome with start positions differing by <1% of the chromosome length, leaving 178 genes. The relationship between expression read count and mean methylation per gene was tested using a negative binomial mixed effects model with glmTBB v.1.1.13^74^ (n=18 hatchlings), including an interaction by feature type given its known influence on gene expression and methylation patterns^75^. Gene ID was also included as a random effect with zero-inflation correction. Significance was assessed via an ANOVA with Type II Wald chi-squared test statistics.

### CV-Field hatchling sex ratio prediction via methylation-based sex markers

We applied the novel methylation-based sex markers to assign CV-Field hatchling sex and determine sex ratios across the timepoint and depth treatment conditions (n=6 timepoints-treatment combinations). To maximise overlap with US-Inc sex marker sites while maintaining data coherence, we retained CpGs (n=19,125,390) covered in at least 75% of CV-Field hatchlings. Missing values for any hatchlings were imputed (7.97% of sex marker sites) with the median methylation value across CV-Field hatchlings within the same relocation timepoint and depth treatment group, then scaled across all US-Inc and CV-Field hatchlings.

Sex prediction was performed via unsupervised k-means clustering to identify natural methylation groupings while capturing experimental and population variation^76^. The optimal number of k clusters was determined using the average silhouette method in factoextra v.1.0.7^77^. K-means clustering was implemented with ‘kmeans’^57^ and visualised via PCAs. Clusters were assigned to predicted sex based on the expectation that female proportions should increase with temperature across timepoints and be higher in warmer shallow versus cooler deep treatments.

To investigate associations with methylation clustering, we performed PERMANOVA analyses on a Euclidean distance matrix of scaled methylation values at the 743 sex markers for all CV-Field hatchlings, implemented in vegan^63^ with 999 permutations. Following earlier logic, if maternal ID was a significant fixed effect in an initial PERMANOVA, it was included as a stratification factor in a subsequent PERMANOVA testing for fixed effects of k-means cluster identity, relocation timepoint, depth treatment, and two-way interactions with k-means cluster identity on methylation clustering. Hatchling sex ratios were calculated as the mean proportion of females per timepoint and treatment combination (Timepoint 1: n=18 hatchlings per treatment; Timepoints 2 and 3: n=20 hatchlings per treatment).

### CV-Field hatchling sex ratio modelling

For comparison with methylation-based sex ratios, we modelled sex ratios using a well-established nest temperature proxy method implemented in embryogrowth v.10.3^11,13^. For the 34 sub-clutches with temperature series data, we fitted a 4-parameter Schoolfield-Sharpe-Magnuson thermal embryonic growth model via maximum likelihood, with recommended fixed parameters for loggerhead turtles^44,78^. Confidence intervals (95%) were computed using the Metropolis-Hastings method of Markov Chain Monte Carlo simulations with 10,000 iterations. Models described the progression of nest temperature and mean hatchling straight carapace length (notch-to-notch SCL; measured with digital callipers for all or up to 20 hatchlings per sub-clutch) over the incubation period. These were used to delimit (1) the middle third of incubation duration (MTID) weighted by embryo growth rate, and (2) the thermosensitive period (TSP; middle third of total development) weighted by embryo growth rate and the thermal reaction norm of sexualization^13^. Sex ratios were then estimated per sub-clutch by projecting the constant temperature equivalent during these periods onto the constant-temperature TSD pattern available for a Northwest Atlantic loggerhead turtle population in Hobe Sound, Florida, USA^39^, with a TPiv of 29.44°C and transitional range of temperatures of 3.759°C (temperature range producing 5% to 95% of each sex), as the TSD pattern has not been measured for the Cabo Verde population. Sex ratio estimates were averaged across each timepoint-depth treatment (Timepoint 1: n=5 per treatment, Timepoint 2: n=8 per treatment, Timepoint 3: n=4 per treatment).

## DATA AVAILABILITY

Sequencing reads for CV-Field hatchlings from Timepoint 2 are available on the European Nucleotide Archive (ENA) under project accession PRJEB75968. Upon publication, all other sequencing reads will be uploaded to ENA under project accession PRJEB104494. Supporting data will be uploaded to a Dryad repository and bioinformatic scripts will be uploaded to a GitHub repository (https://github.com/eugeniecyen/Article_CarCar_SexMarkers). All data can be made available to reviewers on request.

## DECLARATIONS

### Ethics Approval

All sample collection and experiments adhered to national legislation. In Florida, approval was granted by the Florida Fish and Wildlife Commission Marine Turtle (Permit: FWC-MTP-073) and the Florida Atlantic University Institutional Animal Care and Use Committee (Protocol: A21-31), under authorisation to J.W. In Cabo Verde, approval was provided by the Direção Nacional do Ambiente de Cabo Verde (Permit: 037/DNA/2021), under authorisation to C.E.

### Competing Interests

The authors declare they have no competing interests.

### Funding

The study was primarily funded by Queen Mary University of London, UK Research and Innovation (NERC, NE/V001469/1 and NE/X012077/1) grants to C.E and J.M.M-D, and the London NERC Doctoral Training Partnership studentship to E.C.Y (NERC, NE/S007229/1). Further field support was obtained from the National Geographic (NGS-59158R-19) grant to C.E. The temperature-controlled incubator experiments were funded by the Florida National Save the Sea Turtle Foundation, Florida Atlantic University Nelligan Sea Turtle Research Fund, and personal funds to J.W, as well as the Mote Marine Laboratory Research Experience for Graduate Students and personal funds to G.A.C. DNA extractions of the Florida samples were funded by the University of Massachusetts Amherst and NSF-IOS (1904439) grants to L.M.K.

### Author Contributions

E.C.Y and C.E designed the overall study. G.A.C, I.S-R, and J.W designed the artificial incubation experiment, with input from all authors on sample choice. G.A.C conducted egg collection, the experiment, and blood sampling. J.W performed laparoscopic sex identification. B.P.B performed DNA extractions, and L.M.K and C.E organised sequencing. E.C.Y, C.E, K.F, A.T, I.O.A, and D.N performed the field experiment, with logistical support from S.M.C and J.M.M-D. E.C.Y performed DNA extractions and organised sequencing. E.C.Y performed bioinformatic analyses with contributions from J.O.B, J.D.G, and C.E. E.C.Y performed sex ratio modelling with guidance from B.P.B and C.E. C.A.D facilitated collaborations among authors with support from M.L.S. E.C.Y wrote the manuscript with contributions from CE. All authors contributed to the final version of the manuscript.

## Acknowledgements

The authors thank E. Turla, S. Williamson, S. G. Kuschke, J. Perrault, S. Hirsch, M. D. Anderson, R. Germany, and the turtle specialist staff who helped conduct egg collections and the artificial incubator experiment (Florida, USA). We further thank all staff and volunteers with Project Biodiversity for their support throughout the field experiment (Cabo Verde).

## EXTENDED DATA

**Extended Data Table 1.** PERMANOVA results for methylation clustering associations in US-Inc hatchlings. PERMANOVA analyses with 999 permutations against Bray-Curtis distance matrices of methylation values for US-Inc hatchlings (n=29) at the subset of differentially methylated sites (DMS) between sexes (n=777) and all genome-wide CpGs (n=4,816,224). An initial PERMANOVA tested maternal ID as a fixed effect on methylation clustering. If significant, it was included as a stratification factor in a subsequent PERMANOVA testing for fixed effects of sex, temperature treatment, and their interaction represented by a colon (:) symbol. Stars represent significance levels (***: p<0.001, **: p<0.01, *: p<0.05, .: p<0.1).

| Variable | Degrees of freedom | Sum of squares | R<sup>2</sup> | F-value | p-value | Significance code |
| --- | --- | --- | --- | --- | --- | --- |
| **At DMS (n=777)** |  |  |  |  |  |  |
| Methylation ~ Maternal ID |  |  |  |  |  |  |
| Maternal ID | 1 | 0.0099 | 0.0278 | 0.7716 | 0.503 |  |
| Residual | 27 | 0.3450 | 0.9722 |  |  |  |
| Total | 28 | 0.3549 | 1.0000 |  |  |  |
| Methylation ~ Sex + Temperature + Sex : Temperature |  |  |  |  |  |  |
| Sex | 1 | 0.1636 | 0.4610 | 23.2890 | 0.001 | *** |
| Temperature | 1 | 0.0086 | 0.0242 | 1.2211 | 0.228 |  |
| Sex : Temperature | 1 | 0.0071 | 0.0199 | 1.0055 | 0.375 |  |
| Residual | 25 | 0.1756 | 0.4949 |  |  |  |
| Total | 28 | 0.3549 | 1.0000 |  |  |  |
| **At genome-wide CpGs (n=4,816,224)** |  |  |  |  |  |  |
| Methylation ~ Maternal ID |  |  |  |  |  |  |
| Maternal ID | 1 | 0.0060 | 0.0408 | 1.1483 | 0.004 | ** |
| Residual | 27 | 0.1401 | 0.9592 |  |  |  |
| Total | 28 | 0.1461 | 1.0000 |  |  |  |
| Methylation distance ~ Sex + Temperature + Sex : Temperature (strata: Maternal ID) |  |  |  |  |  |  |
| Sex | 1 | 0.0053 | 0.0361 | 1.0136 | 0.146 |  |
| Temperature | 1 | 0.0055 | 0.0375 | 1.0524 | 0.044 | * |
| Sex : Temperature | 1 | 0.0053 | 0.0360 | 1.0107 | 0.270 |  |
| Residual | 25 | 0.1301 | 0.8904 |  |  |  |
| Total | 28 | 0.1461 | 1.0000 |  |  |  |

**Extended Data Table 2.** Genes associated with differentially methylated sites in the top 5% outlier subset. Genes associated with differentially methylated sites (DMS) identified as top 5% outliers in relation to absolute mean methylation difference between the sexes of US- Inc hatchlings (n=17 DMS associated with 14 genes). Starred DMS (*) were associated with genes that had direct roles in sex development or gametogenesis (**Supplementary Table 3**). See **Supplementary Table 1** for all 29 outlier DMS, including those not associated with annotated genes.

| DMS ID | Chromosome | Site | Feature type | Methylation difference (%) | q-value | Gene name |
| --- | --- | --- | --- | --- | --- | --- |
| **Methylated in females** |  |  |  |  |  |  |
| TRIT1.3 | 19 | 7889086 | Intron | 37.4 | 0.00031 | tRNA Isopentenyltransferase 1 |
| TRIT1.2 | 19 | 7889078 | Intron | 35.3 | 0.00001 | tRNA Isopentenyltransferase 1 |
| AMDHD2.1* | 10 | 48194734 | Promoter | 34.5 | 0.00663 | Amidohydrolase Domain Containing 2 |
| CTNNA2.1 | 4 | 134907444 | Intron | 33.2 | 0.00117 | Catenin Alpha 2 |
| DIP2C.1 | 2 | 23069168 | Intron | 32.6 | 0.00013 | Disco-interacting Protein 2 Homolog C |
| RASGRP3.1 | 3 | 62872080 | Intron | 32.5 | 0.00008 | RAS Guanyl Releasing Protein 3 |
| EHHADH.1 | 9 | 49010864 | Intron | 30.0 | 0.00038 | Enoyl-CoA Hydratase And 3-Hydroxyacyl CoA Dehydrogenase |
| NCKAP5.1 | 11 | 63225940 | Exon | 29.4 | 0.00004 | NCK Associated Protein 5 |
| TRIT1.4 | 19 | 7889097 | Intron | 28.5 | 0.04759 | tRNA Isopentenyltransferase 1 |
| PPM1E.1 | 17 | 20633049 | Intron | 27.9 | 0.04932 | Protein Phosphatase, Mg2+/Mn2+ Dependent 1E |
| SNTG2.2 | 3 | 163195550 | Intron | 27.8 | 0.00101 | Syntrophin Gamma 2 |
| TRIT1.1 | 19 | 7889065 | Intron | 27.6 | 0.02749 | tRNA Isopentenyltransferase 1 |
| **Methylated in males** |  |  |  |  |  |  |
| PCDH10.1 | 4 | 52370405 | Intergenic | 32.9 | 0.00464 | Protocadherin 10 |
| RTN1.1 | 6 | 112795770 | Intron | 29.2 | 0.00087 | Reticulon 1 |
| FAM110C.1 | 1 | 250973500 | Intron | 28.8 | 0.00371 | Family With Sequence Similarity 110 Member C |
| RPL13.1 | 17 | 8464636 | Intergenic | 28.0 | 0.00216 | Ribosomal Protein L13 |
| HSP90AB1.1* | 3 | 61514327 | Intergenic | 27.8 | 0.00844 | Heat Shock Protein 90 kDa Alpha Class B Member 1 |

**Extended Data Table 3.**
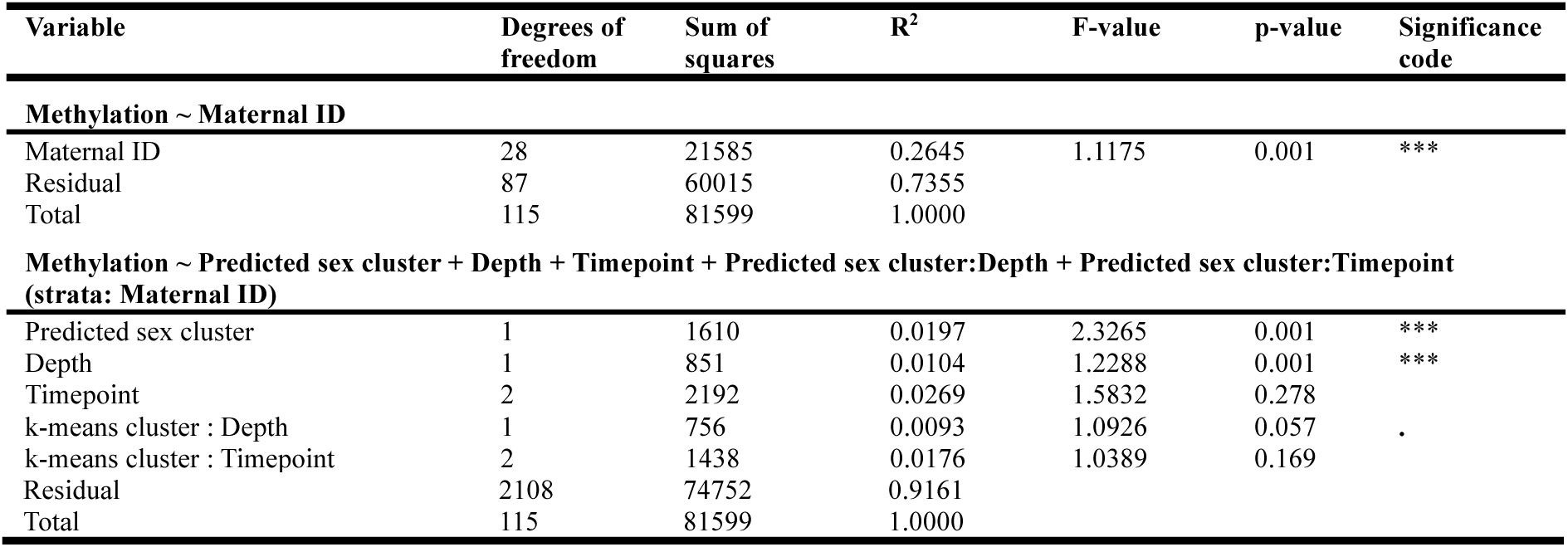
PERMANOVA results for methylation clustering associations in CV-Field hatchlings. PERMANOVA analyses with 999 permutations against a Euclidean distance matrix of scaled methylation values for CV-Inc hatchlings (n=116) at present methylation-based sex marker sites (n=743). An initial PERMANOVA tested maternal ID as a fixed effect on methylation clustering. This was significant and thus included as a stratification factor in a subsequent PERMANOVA testing for fixed effects of predicted sex cluster identity, depth treatment, relocation timepoint, and all two-way interactions with predicted sex cluster identity. Stars represent significance levels (***: p<0.001, **: p<0.01, *: p<0.05, .: p<0.1).

| Variable | Degrees of freedom | Sum of squares | R<sup>2</sup> | F-value | p-value | Significance code |
| --- | --- | --- | --- | --- | --- | --- |
| **Methylation ~ Maternal ID** |  |  |  |  |  |  |
| Maternal ID | 28 | 21585 | 0.2645 | 1.1175 | 0.001 | *** |
| Residual | 87 | 60015 | 0.7355 |  |  |  |
| Total | 115 | 81599 | 1.0000 |  |  |  |
| **Methylation ~ Predicted sex cluster + Depth + Timepoint + Predicted sex cluster:Depth + Predicted sex cluster:Timepoint (strata: Maternal ID)** |  |  |  |  |  |  |
| Predicted sex cluster | 1 | 1610 | 0.0197 | 2.3265 | 0.001 | *** |
| Depth | 1 | 851 | 0.0104 | 1.2288 | 0.001 | *** |
| Timepoint | 2 | 2192 | 0.0269 | 1.5832 | 0.278 |  |
| k-means cluster : Depth | 1 | 756 | 0.0093 | 1.0926 | 0.057 | . |
| k-means cluster : Timepoint | 2 | 1438 | 0.0176 | 1.0389 | 0.169 |  |

|  |  |  |  |
| --- | --- | --- | --- |
| Residual | 2108 | 74752 | 0.9161 |
| Total | 115 | 81599 | 1.0000 |

**Extended Data Fig. 1.**
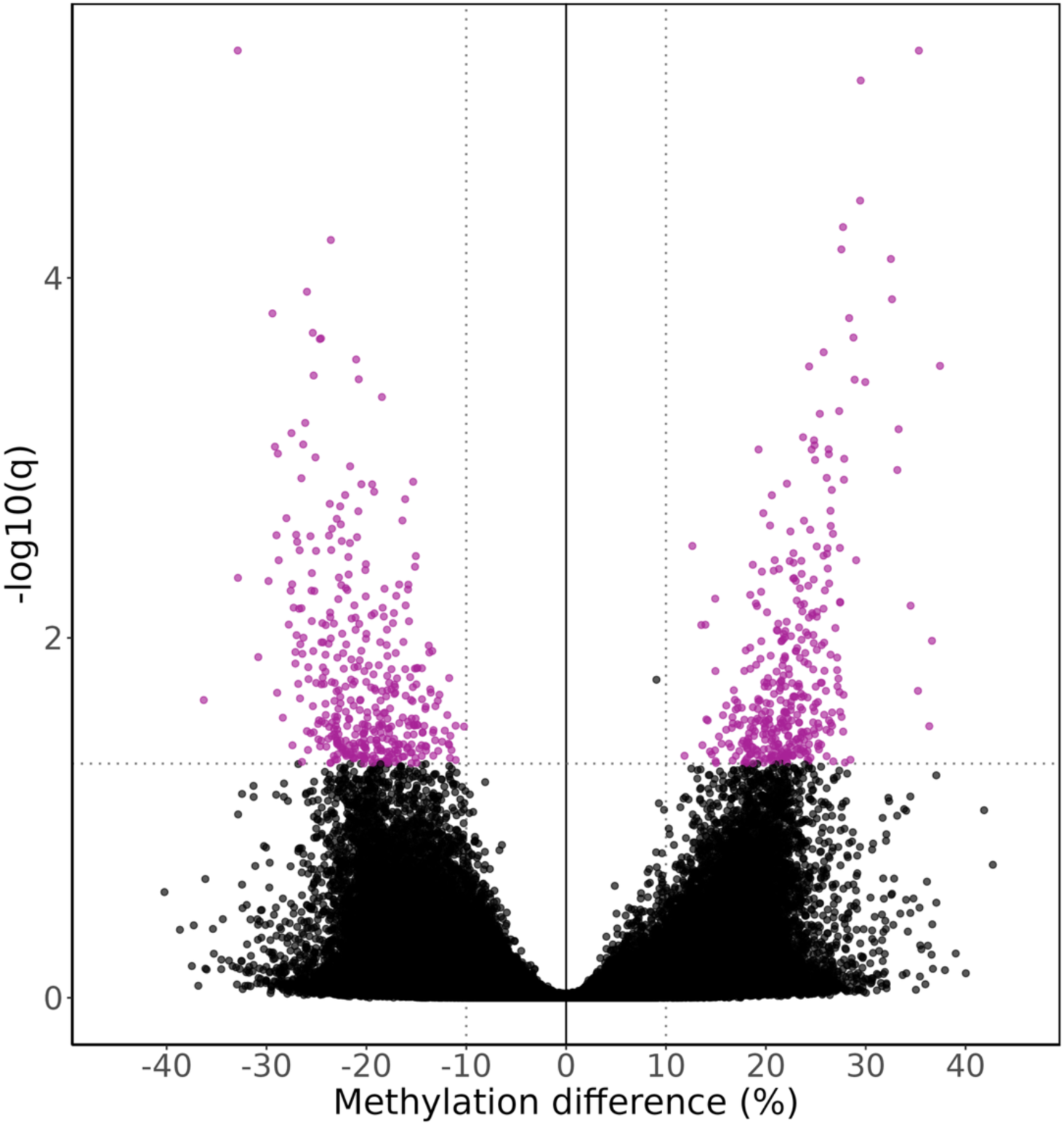
Volcano plot of methylation differences between the sexes of US- Inc hatchlings. Mean methylation difference (%) between the sexes against the -log10(q-value) for all genome-wide CpGs (n=4,186,224 total), with differentially methylated sites (DMS, n=777) in purple. Dotted lines represent the thresholds used to classify a CpG site as differentially methylated between the sexes (mean methylation difference > 10%, q<0.05).

**Extended Data Fig. 2.**
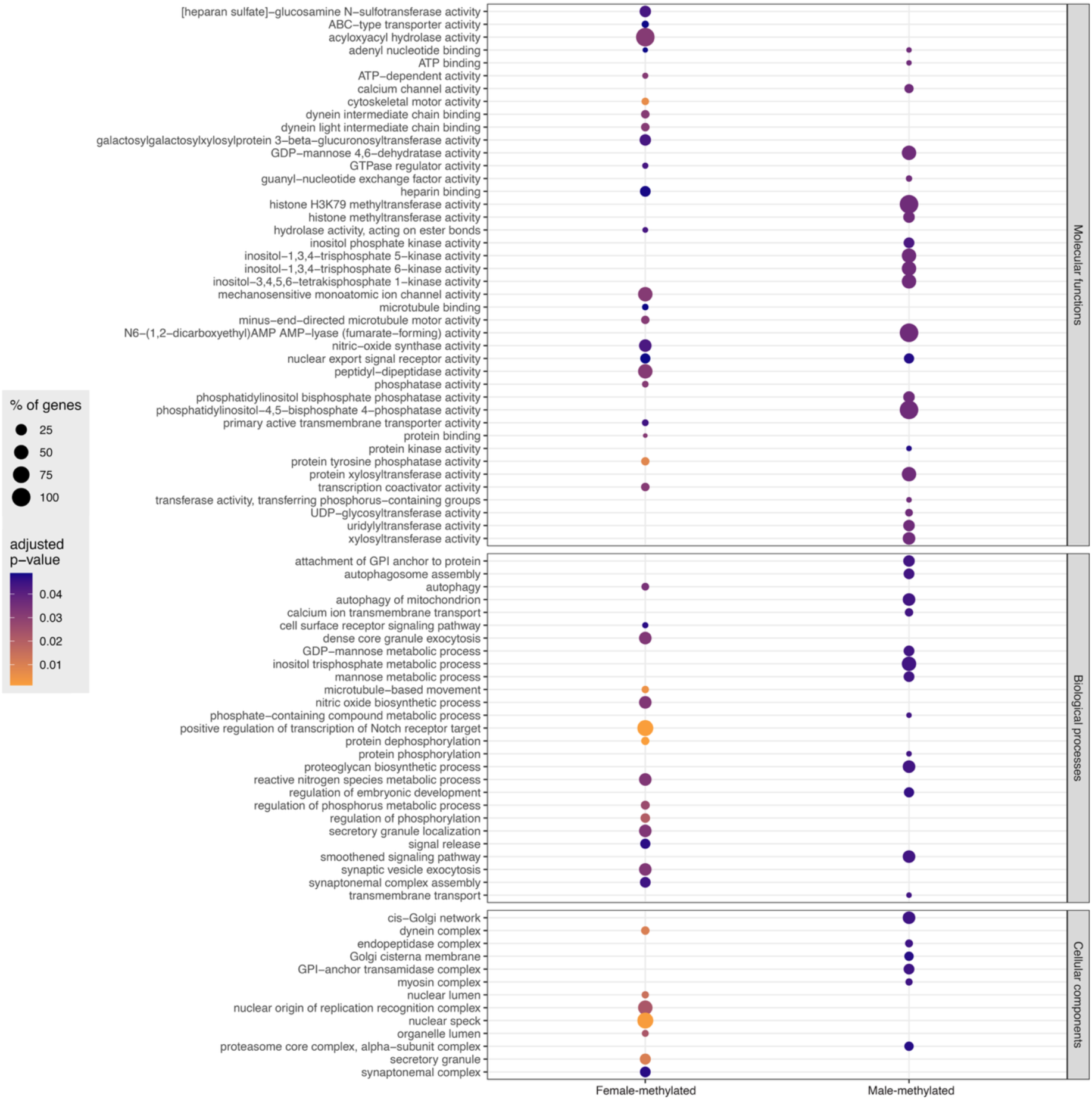
Gene Ontology enrichment analysis of sex-specific methylation in US-Inc hatchlings. Gene Ontology terms enriched for genes associated to differentially methylated sites (DMS) between the sexes of US-Inc hatchlings (n=44 enriched terms for female-hypermethylated genes; n=40 enriched terms for male-hypermethylated genes; n=84 total). Dot size represents the percentage of genes for a given term enriched in the dataset, coloured by the adjusted p-value.

**Extended Data Fig. 3.**
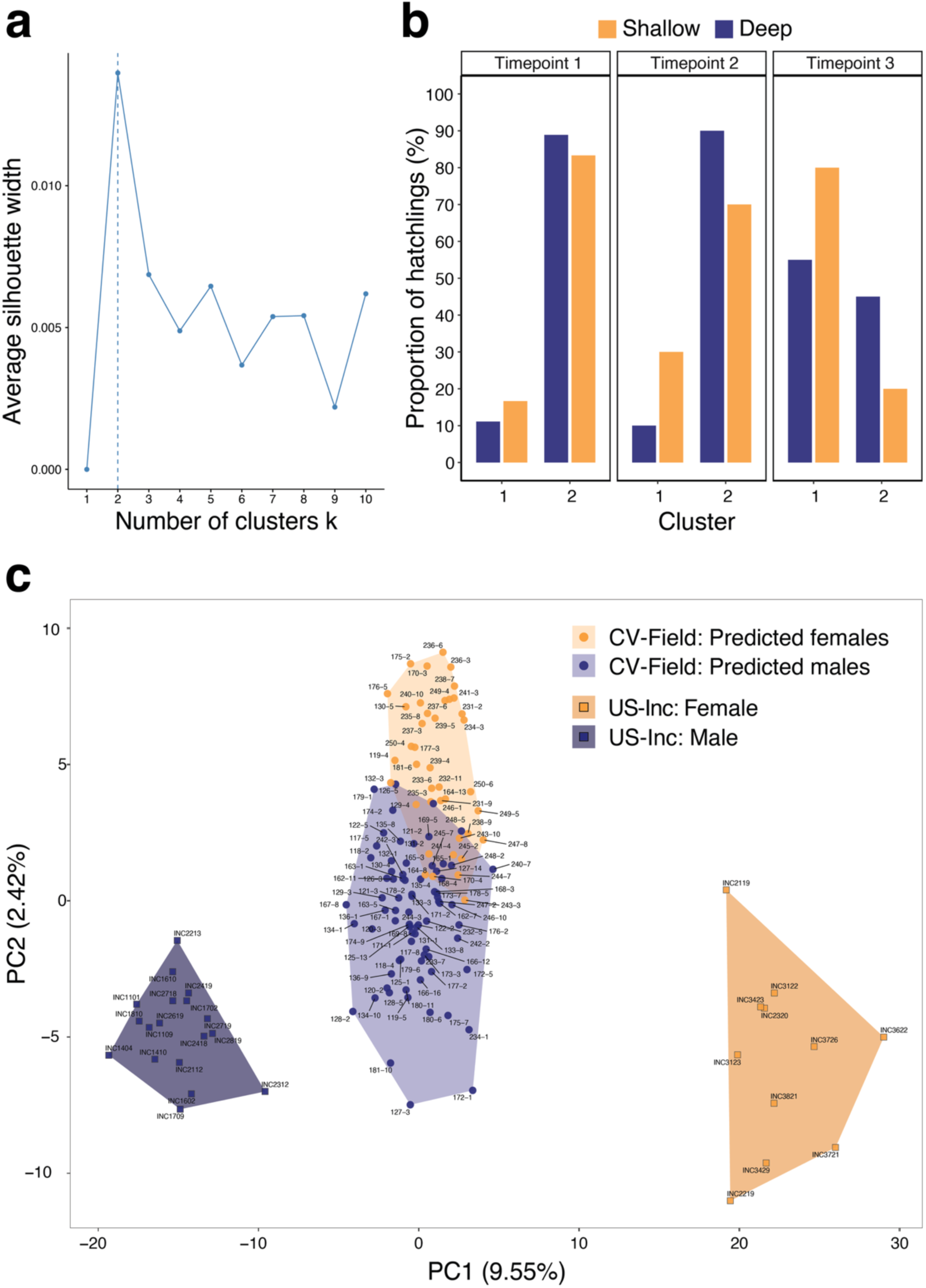
Accessory plots for methylation-based sex prediction of CV-Field hatchlings. **(a)** The number of optimal k-means clusters identified via the average silhouette method from methylation variance at 743 methylation-based sex markers of the CV-Field hatchlings (n=116). **(b)** Proportion of CV-Field hatchlings belonging to each k-means cluster, calculated across all hatchlings per timepoint-depth treatment (Timepoint 1: n=18 hatchlings per treatment; Timepoints 2 and 3; n=20 hatchlings per treatment; n=116 total). Colours represent depth treatment (shallow: yellow, deep: blue). **(c)** Principal component analysis of all hatchlings (n=29 US-Inc hatchlings; n=116 CV-Field hatchlings; n=145 total) at 743 methylation-based sex markers and overlap between both datasets. The proportion of methylation variance explained by each principal component (PC) is shown in brackets on each axis. US-Inc hatchlings are represented by squares and coloured by sex (females: darker yellow, males: darker blue) and CV-Field hatchlings are represented by circles and coloured by the predicted sex cluster (predicted females: paler yellow, predicted males: paler blue).

**Extended Data Fig.4.**
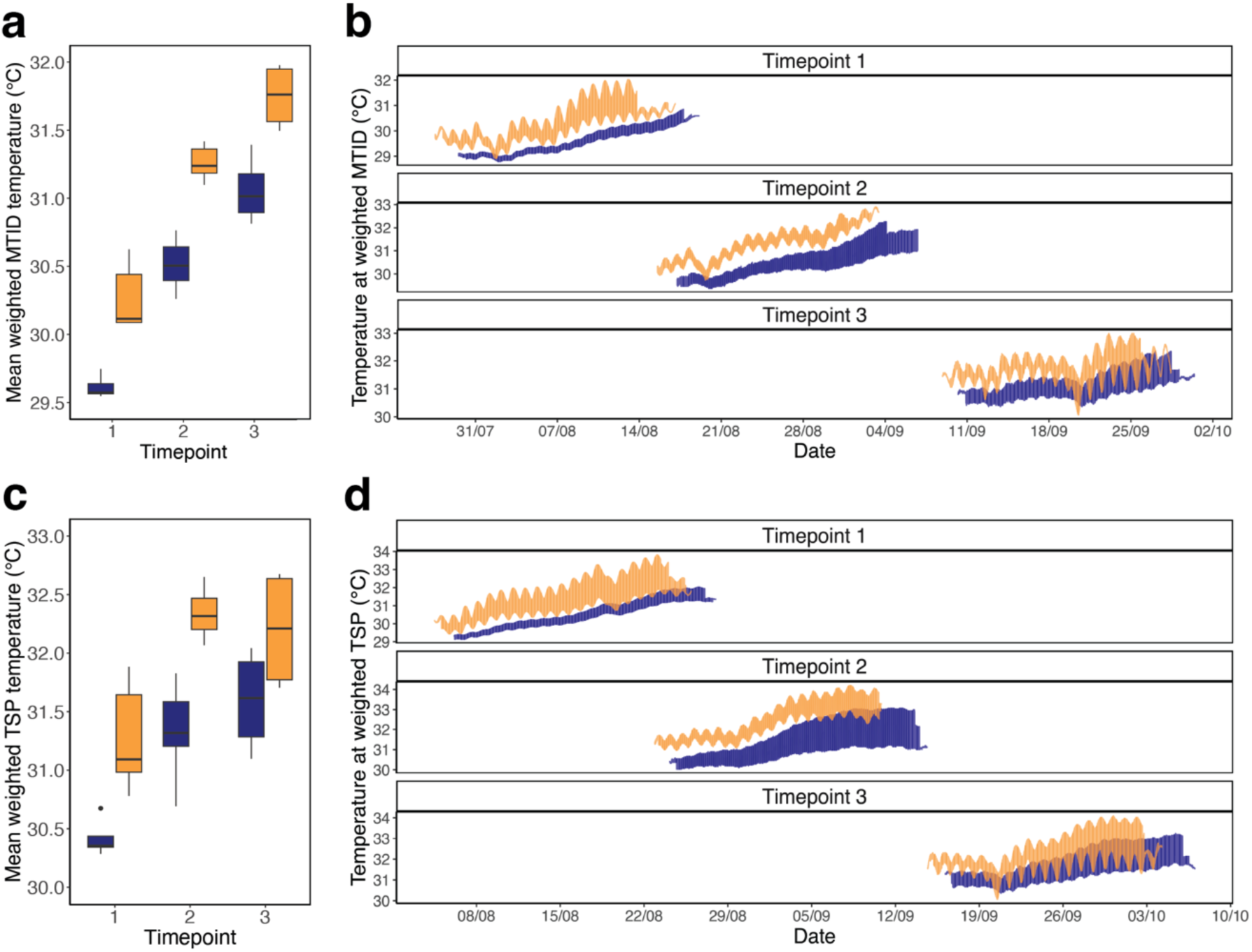
Nest temperature modelling results. Constant temperature equivalents derived from a thermal embryonic growth model for sub-clutches with temperature data per timepoint and depth treatment (Timepoint 1: n=5 per treatment, Timepoint 2: n=8 per treatment, Timepoint 3: n=4 per treatment; n=34 total). **(a-b)** Temperatures (°C) modelled during the weighted middle third of incubation (MTID), with the **(a)** mean temperature and **(b)** temperature series shown across all sub-clutches per timepoint and depth treatment. **(c-d)** Temperatures (°C) modelled during the weighted thermosensitive period (TSP), with the **(a)** mean temperature and **(b)** temperature series shown across all sub-clutches per timepoint and depth treatment. All plots are coloured by depth treatment (shallow: yellow, deep: blue). For all boxplots, the central line represents the median, the box depicts the interquartile range (IQR), and the whiskers represent 1.5 times the IQR.

**Extended Data Fig.5.**
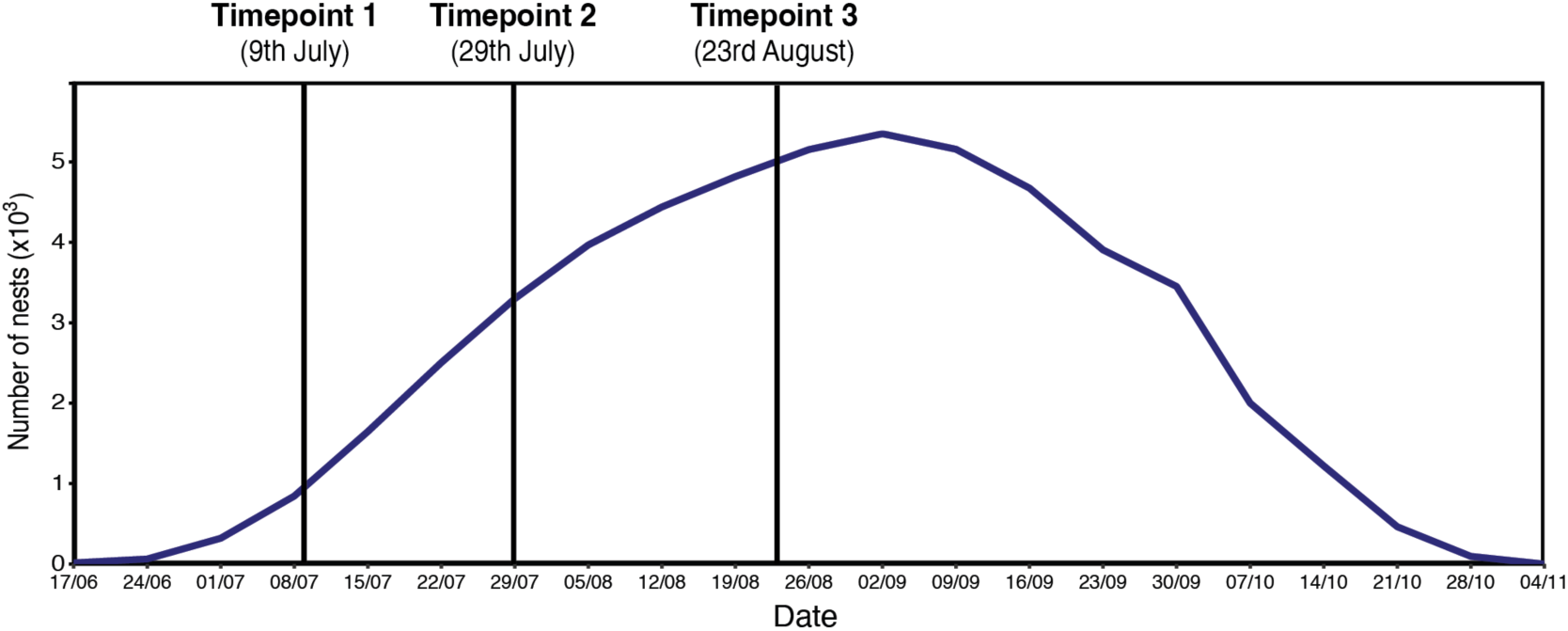
Nesting numbers over the 2021 season. Total nest numbers (n=53,498) laid throughout the 2021 nesting season in Sal, Cabo Verde, censused by Project Biodiversity. Vertical bars indicate the clutch relocation timepoint in the field experiment.

## SUPPLEMENTARY INFORMATION

**Supplementary Table 1. Functional annotations of all gene-associated DMS in US-Inc hatchlings.** Functional information for all differentially methylated sites (DMS) between sexes of US-Inc hatchlings associated with an annotated gene (n=436 DMS, associated with 392 genes).

**Supplementary Table 2. Gene Ontology terms linked to DMS-associated genes in US-Inc hatchlings.** Gene Ontology terms enriched for genes associated to differentially methylated sites (DMS) between sexes of US-Inc hatchlings (n=44 for female-hypermethylated genes; n=40 for male-hypermethylated genes; n=84 total).

**Supplementary Table 3. DMS-associated genes with sex development-related roles in US- Inc hatchlings.** Genes associated with differentially methylated sites (DMS) between sexes of US-Inc hatchlings, with direct roles in sex development or gametogenesis pathways reported in the primary literature (n=40 genes, associated to 46 DMS). For genes with multiple DMS, the mean methylation difference between sexes (%) was calculated across all DMS identified on that gene. Starred genes (*) also possessed DMS within the top 5% methylation outlier subset overall (**Extended Data Table 2**).

**Supplementary Table 4. Nest temperature and sex ratio modelling results.** Outputs from a thermal embryonic growth model fit to sub-clutches with recorded temperature series from the Northeast Atlantic field experiment (Timepoint 1: n=5 per depth treatment, Timepoint 2: n=8 per depth treatment, Timepoint 3: n=4 per depth treatment; n=34 total). Constant temperature equivalents were modelled during the weighted middle third of incubation (MTID) and the weighted thermosensitive period (TSP). Sex ratio per sub-clutch was estimated by applying a constant-temperature TSD pattern with a pivotal temperature of 29.44°C.

**Supplementary Table 5. Metadata and methylation statistics for US-Inc hatchlings.** Sample metadata and methylation summary statistics for US-Inc hatchlings from the Northwest laboratory incubation experiment (n=29).

**Supplementary Table 6. Metadata and methylation statistics for CV-Field hatchlings.** Sample metadata and methylation summary statistics for CV-Field hatchlings from the Northeast Atlantic field experiment (n=116).

## Notes

### Competing Interest Statement

The authors have declared no competing interest.

